# Alterations in gut microbiome composition and function in irritable bowel syndrome and increased probiotic abundance with daily supplementation

**DOI:** 10.1101/2021.09.02.458777

**Authors:** Joann Phan, Divya Nair, Suneer Jain, Thibaut Montagne, Demi Valeria Flores, Andre Nguyen, Summer Dietsche, Saurabh Gombar, Philip Cotter

**Affiliations:** Sun Genomics, Inc., San Diego, CA, USA; Atropos Health, Palo Alto, CA, USA; ResearchDx, Inc., Irvine, CA, USA

**Author notes:** **Corresponding Author:** Joann Phan, **Email:**.

**Keywords:** irritable bowel syndrome, gut microbiome, metagenomics, probiotics

## Abstract

**Background:** Irritable bowel syndrome (IBS) is characterized by abdominal discomfort and irregular bowel movements and stool consistency. Because there are different symptoms associated with IBS, it is difficult to diagnose the role of the microbiome in IBS.

**Objective:** Here, we present a study that includes metagenomic sequencing of stool samples from subjects with the predominant subtypes of IBS and a healthy cohort. We collected longitudinal samples from individuals with IBS who took daily made-to-order precision probiotic and prebiotic supplementation throughout the study.

**Materials and Methods:** This study includes a population of 489 individuals with IBS and 122 healthy controls. All stool samples were subjected to shotgun metagenomic sequencing. Precision probiotics and prebiotics were formulated for all subjects with longitudinal timepoints.

**Results:** There was significant variation explained in the microbiome between the healthy and IBS cohorts. Individuals with IBS had a lower gut microbiome diversity and reduced anti-inflammatory microbes compared to the healthy controls. *Eubacterium rectale* and *Faecalibacterium prausnitzii* were associated with healthy microbiomes while *Shigella* species were associated with IBS. Pathway analysis indicated a functional imbalance of short chain fatty acids, vitamins, and a microbial component of Gram-negative bacteria in IBS compared to healthy controls. In the longitudinal dataset, there was a significant difference in microbiome composition between timepoints 1 and 3. There was also a significant increase in the overall microbiome score and relative abundances of probiotic species used to target the symptoms associated with IBS.

**Conclusions:** We identified microbes and pathways that differentiate healthy and IBS microbiomes. In response to precision probiotic supplementation, we identified a significant improvement in the overall microbiome score in individuals with IBS. These results suggest an important role for probiotics in managing IBS symptoms and modulation of the microbiome as a potential management strategy.

**Importance:** An estimated 35 million people in the United States and 11.5% of the population globally are affected by IBS. Immunity, genetics, environment, diet, small intestinal bacterial overgrowth (SIBO), and the gut microbiome are all factors that contribute to the onset or triggers of IBS. With strong supporting evidence that the gut microbiome may influence symptoms associated with IBS, elucidating the important microbes that contribute to the symptoms and severity is important to make decisions for targeted treatment. As probiotics have become more common in treating IBS symptoms, identifying effective probiotics may help inform future studies and treatment.

## Introduction

Irritable bowel syndrome (IBS) is characterized by chronic gastrointestinal discomfort and abdominal pain with changes in bowel habits or stool consistency. IBS affects approximately 11.5% of the population, depending on the country or region (1). Because of the high prevalence of IBS, symptoms contribute to changes in quality of life and increases in healthcare and economic burden (2–5). There are four subtypes based on the symptoms people experience, IBS-C (constipation), IBS-D (diarrhea), IBS-A (alternating), or unspecified (6). Individuals with IBS-A experience alternating symptoms of chronic diarrhea and constipation. The criterion for diagnosis is symptom based and codified in the Rome IV criteria; there is not yet consensus on the underlying etiology of IBS (7, 8). In addition, there are different factors that contribute to the varying symptoms of IBS, including diet, immune response, host genetics, environmental stress, gut microbiome composition, and dysbiosis (9, 10).

Currently, the role of the gut microbiome in individuals with IBS remains poorly understood. A “healthy” gut microbiome may be undefined, but there are microorganisms associated with an unhealthy microbiome, including microorganisms that induce inflammation or dysbiosis that contribute to the symptoms associated with IBS. Changes in microbiome composition also impact the microbiome’s functional potential and metabolism, which may in turn affect host physiology. For example, studies indicate individuals experiencing IBS-C show microbiome signatures such as increased *Pseudomonas* and *Bacteroides thetaiotamicron* with depletion of *Paraprevotella*, significant associations with *Fusobacterium nucleatum* and *Meganomoas hypermegale*, and pathways of sugar and amino acid metabolism (11). In addition, research has characterized the microbiome of subjects with IBS-C with the biosynthetic pathways for sugar and amino acid metabolism, subjects with IBS-D had microbes that predominated the pathways for nucleotides and fatty acids acid synthesis. (11). Amplicon studies have also described an enrichment of Clostridiales, *Prevotella*, and Enterobacteriaceae, reduced microbial richness, and the presence of methanogens in IBS (12, 13). However, amplicon studies can be subject to amplification bias, yielding variable results and do not resolve species-level taxonomic classification. Alternatively, several studies limited by sample size and methodology have not shown a difference between a healthy cohort and individuals with IBS (14).

Because of the differences in IBS symptoms people experience and the individual nature of the syndrome, there is no standardized treatment or dietary recommendations to alleviate IBS symptoms (15). The antibiotic rifaximin has been shown to be an effective treatment for IBS-D (16, 17). However, rifaximin is ineffective for all IBS subtypes and antibiotic usage may be associated with an increased risk for IBS (18–21). There are additional options for treatment, including pharmaceutical options and fecal transplants, but these options are not always feasible and can be invasive. The administration of live microbial organisms, in the form of probiotics, has gained popularity with patients to alleviate their symptoms. Probiotics can alter the microbiome of patients with and without IBS (22, 23), depending on their endogenous microbiomes (24). Microbes not present in the current gut microbiome can also be re-established through probiotic supplementation (24). In individuals with IBS, there is correlative depletion of *Bifidobacterium and Lactobacillus* (8). Therefore, re-introducing probiotics into the gut microbiomes of individuals with IBS may lead to phenotypic changes. Clinical trials have demonstrated the reduction of symptoms associated with IBS with probiotic supplementation. Further studies have shown that in subjects with IBS-D and treated with *Bifidobacterium longum, B. bifidum, B. lactis, B. infantis*, and *Lactobacillus acidophilus*, there is a change in inflammation-related metabolites (25). Individuals with IBS on a gluten-free diet with probiotic supplementation of *Lactobacillus* and *Bifidobacterium spp*. saw an overall improvement in symptoms (26). Probiotic supplementation has also reduced stomach pain and improved stool consistency in individuals with IBS (27, 28).

Here, we present a large-scale metagenomic study to characterize and compare the microbiome composition and functional potential of healthy controls and individuals with IBS. In addition, we collected longitudinal timepoints from individuals with IBS on daily prebiotic and probiotic supplementation. Our primary goals were to 1) identify metagenomic signatures associated with IBS and 2) investigate the microbiome effects of precision prebiotics and probiotics on individuals with IBS. Each made-to-order formulation includes 4-8 probiotic strains and 1-3 prebiotics, each at different concentrations from a biobank of over 100 possible ingredients supported by the clinical literature. Whole genome shotgun sequencing allows for species-level resolution and identification of functional potential. This method reduces amplification bias and allows for sequence-based mapping of pathways rather than functional prediction based on amplicon-based taxonomic classification. We also investigate whether traditional tools in gross microbiome analysis can determine changes to the microbiome after probiotic supplementation. We hypothesized that metagenomic features distinguish healthy vs. IBS microbiome subtypes and that daily probiotic supplementation modulates the microbiomes of the individuals with IBS.

## Results

### IBS and healthy subject demographics

We included a total of 611 subjects in this study. Subjects without reported comorbidities and self-reported as healthy were included as the healthy control population. There were 489 subjects with IBS and 122 subjects in the healthy control population (Table 1). In addition, longitudinal samples from people with IBS were assessed to identify specific microbiome changes during the course of prebiotic and probiotic supplementation. These healthy and IBS subjects were also assigned an internal health index score for their initial microbiome profile and subsequent timepoints. The rationally designed and scientifically backed probiotics were part of a daily regimen for all subjects. Longitudinal timepoints were approximately 4 months apart. Of the 489 IBS subject population, 134 subjects had at least 2 timepoints, 56 subjects had 3 timepoints, 28 subjects with 4 timepoints, 15 subjects with 5 timepoints, 5 subjects with 6 timepoints, and 1 subject with 7 timepoints.

**Table 1.**
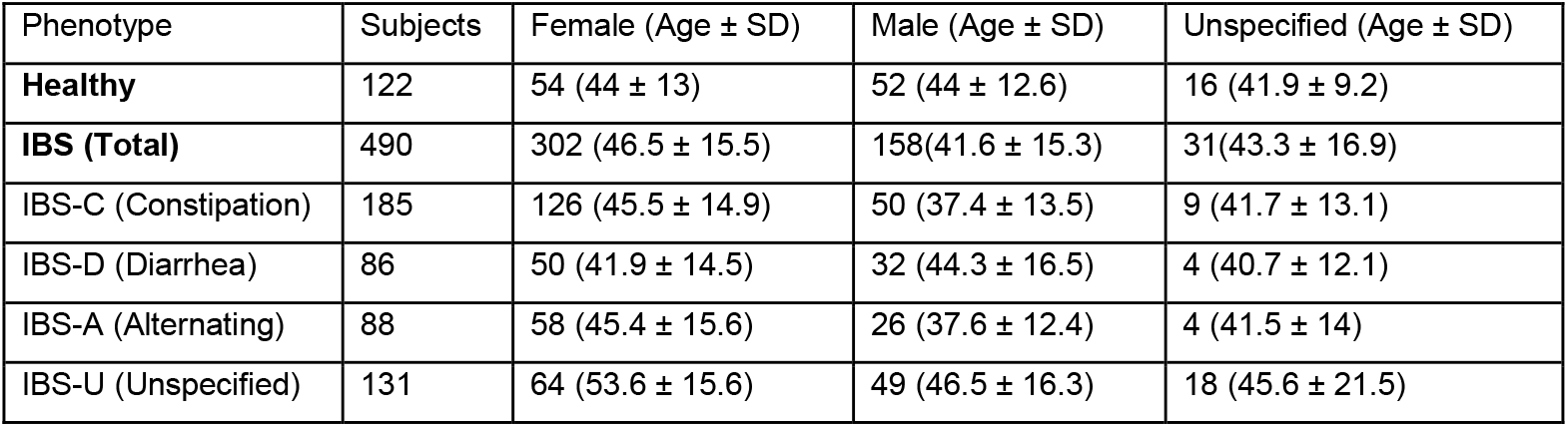
Subject demographics. “Healthy” controls are self-reported as healthy subjects with no existing comorbidities. Subjects with IBS are also self-reported. The subtype designation is based on the subject symptoms. For the alternating designation, subjects experienced symptoms of constipation and diarrhea. Cohort populations are further classified by gender. Average and standard deviation of age groups are listed next to each population.

| Phenotype | Subjects | Female (Age ± SD) | Male (Age ± SD) | Unspecified (Age ± SD) |
| --- | --- | --- | --- | --- |
| **Healthy** | 122 | 54 (44 ± 13) | 52 (44 ± 12.6) | 16 (41.9 ± 9.2) |
| **IBS (Total)** | 490 | 302 (46.5 ± 15.5) | 158(41.6 ± 15.3) | 31(43.3 ± 16.9) |
| IBS-C (Constipation) | 185 | 126 (45.5 ± 14.9) | 50 (37.4 ± 13.5) | 9 (41.7 ± 13.1) |
| IBS-D (Diarrhea) | 86 | 50 (41.9 ± 14.5) | 32 (44.3 ± 16.5) | 4 (40.7 ± 12.1) |
| IBS-A (Alternating) | 88 | 58 (45.4 ± 15.6) | 26 (37.6 ± 12.4) | 4 (41.5 ± 14) |
| IBS-U (Unspecified) | 131 | 64 (53.6 ± 15.6) | 49 (46.5 ± 16.3) | 18 (45.6 ± 21.5) |

### Reduced microbial diversity and microbial signatures associated with IBS

First, to compare the microbial community composition between the IBS and healthy control populations, a principal coordinates analysis was performed to visualize the beta diversity between the two cohorts (Figure 1). All healthy control microbiome samples clustered tightly together, while there was a spread of IBS samples that clustered around and away from the healthy control microbiome samples. The differences in phenotypes were identified with increased relative abundances of Enterobacterales species and reduction in *Eubacterium rectale* and *Faecalibacterium prausnitzii* (Figure 1). A subset of microbes that distinguish healthy and IBS were determined by random forest and were plotted along the second principal coordinate axis to show the spread of sample clustering between the healthy and IBS microbiomes. Next, when calculating alpha diversity metrics, there was a significant reduction in the Shannon index, richness, and evenness in IBS subtypes compared to the healthy control population (Figure 1).

**Figure 1.**
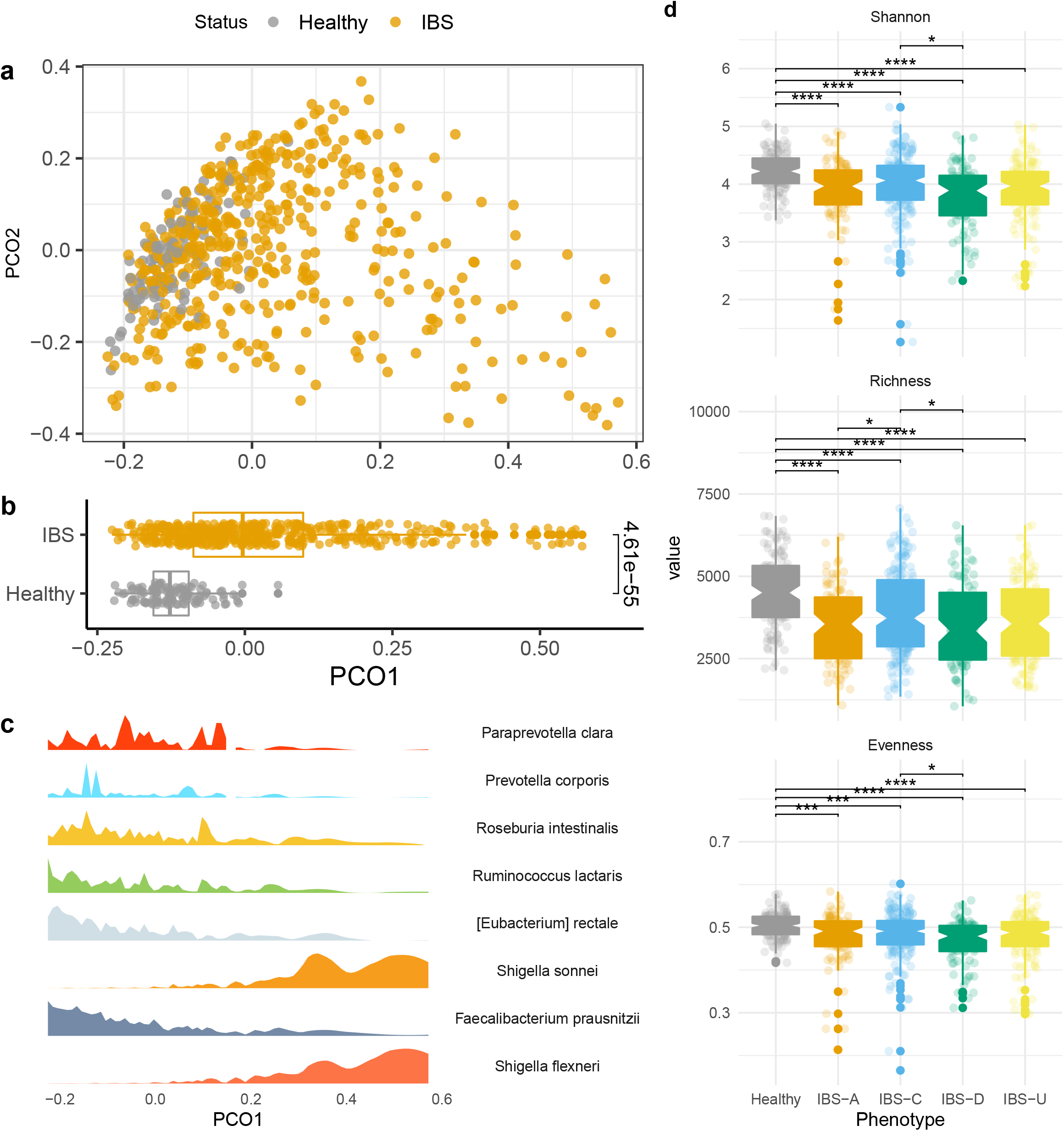
Microbiome profiles of the healthy and IBS cohorts. A) Principal coordinates analysis based on the Bray-Curtis dissimilarity distance matrix of the IBS and healthy microbiomes. B) Boxplot of the microbiome distributions along the PCO1 axis. An unpaired t-test was computed. C) A random forest was employed to differentiate microbes between healthy and IBS subtypes. The density of microbes selected from random forest correspond to the sample distribution along PCO1 axis. D) Alpha diversity between healthy and each IBS subtype cohort. The Shannon index, species richness, and evenness were calculated. Unpaired t-tests were conducted, and p-values were adjusted with Benjamin-Hochberg false discovery rate (FDR) for multiple comparisons. * p < 0.05, ** p < 0.01, *** p < 0.001, **** p < 0.0001.

Based on whole-genome shotgun metagenomic sequencing, microbial signatures distinguish the healthy control and IBS populations. Using a permutated multivariate analysis of variance, we calculated a significant variation that explained the difference between the microbiome of healthy and IBS subtypes (R2 = 0.028, p < 0.001). We performed a random forest analysis to identify the distinguishing microbes between healthy and IBS phenotypes. To identify statistically significant changes in the relative abundances of microbes within healthy or IBS subtypes, we performed an unpaired t-test and adjusted p-values for multiple testing corrections. This analysis revealed *Eubacterium rectale* and *Faecalibacterium prausnitzii* as significantly increased microbial species in the healthy control population relative to all IBS subtypes (Figure 2), while we found inflammatory species of *Shigella* elevated in IBS (Figure 2). We further interrogated the microbial differences between IBS subtypes and found that *Paraprevotella clara, Prevotella corporis, Roseburia intestinalis* and *Ruminococcus lactaris* significantly decreased in different IBS subtypes relative to the healthy control population (Figure 2).

**Figure 2.**
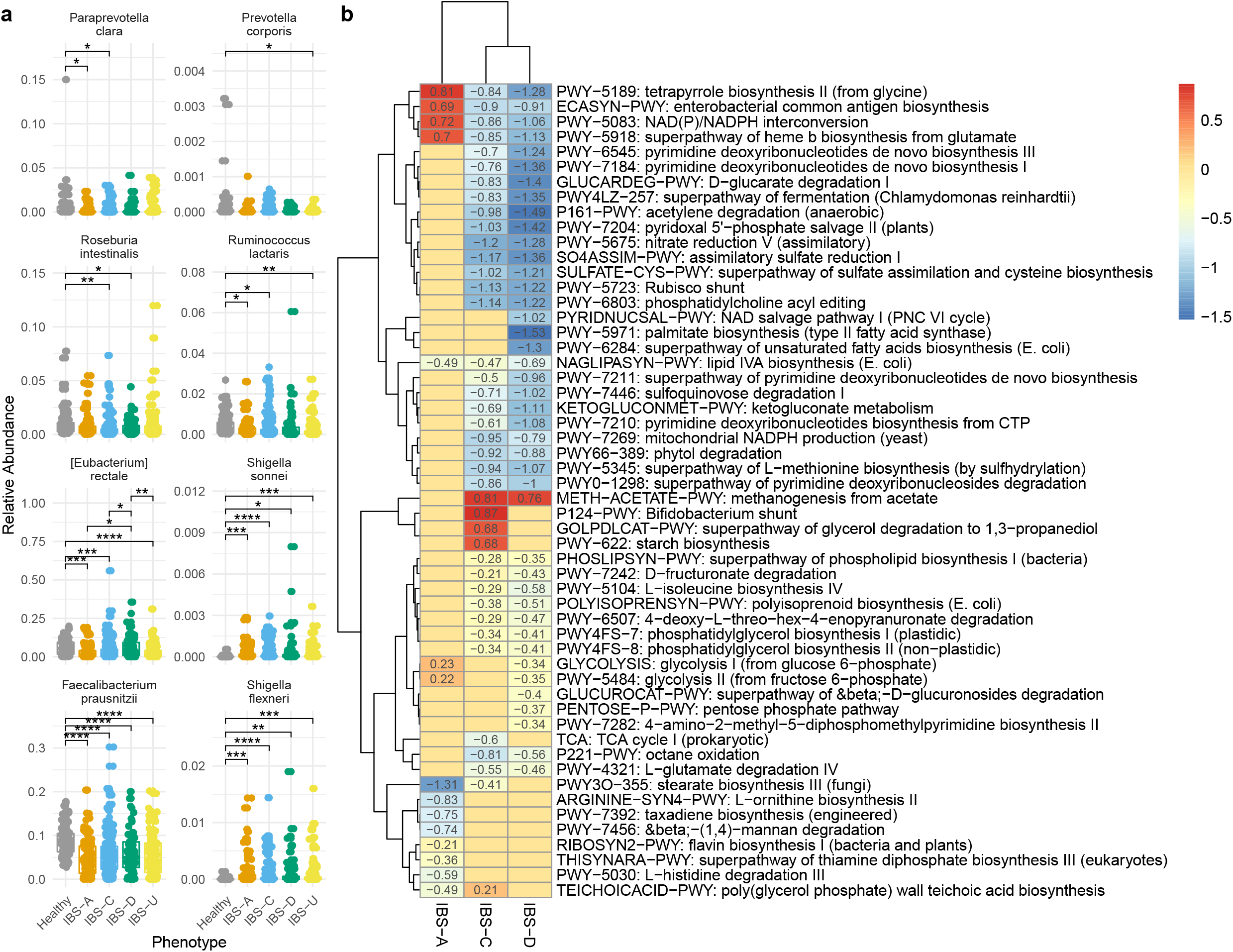
Microbes and pathways that differentiate healthy and IBS cohorts. A) Relative abundances of the microbes associated with healthy and IBS populations. A random forest was used to determine microbes that contribute to differentiating healthy and IBS. The relative abundances of a subset of the microbes were plotted for healthy and each IBS subtype. T-test were calculated. P values were adjusted for multiple comparisons testing by false discovery rate corrections. Non-significant comparisons were omitted. B) Functional pathways associated with healthy and IBS gut microbiomes. Multivariate linear association testing with Maaslin2 was used to determine pathways associated with IBS relative to the healthy control population. Values indicate the beta coefficient from linear association testing. Pathways listed were filtered based on q value < 0.1 and beta coefficients > 0.2 or < −0.2. * p < 0.05, ** p < 0.01, *** p < 0.001, **** p < 0.0001.

### Functional profile of the gut microbiome of subjects with IBS and healthy

To determine the functional profiles of the gut microbiome associated with IBS, we mapped the metagenomic reads against the MetaCyc database with Humann3 to identify pathway abundances. There was a total of 471 pathways detected in the metagenomes of healthy and individuals with IBS. Multivariate linear association testing identified pathways associated with each IBS dominant subtype relative to the healthy control cohort (Figure 2). Pathways involved in tetrapyrrole biosynthesis from glycine, enterobacterial common antigen biosynthesis, NADP/NADPH interconversion, and the super pathway of heme b biosynthesis from glutamate were positively associated with IBS-A (Figure 2). Methanogenesis from acetate was associated with IBS-C and IBS-D (Figure 2). Pathways involved in the *Bifidobacterium* shunt, the super pathway of glycerol degradation to 1,3-propanediol, and starch biosynthesis were associated with IBS-C (Figure 2). Meanwhile, pathways associated with amino acid and ribonucleotide biosynthesis, polysaccharide degradation, and fermentation were associated with healthy microbiome functional profiles (Figure 2).

### Probiotics may modulate the microbiome of subjects with IBS

Within a subset of the IBS population, there were 134 individuals with at least two timepoints and 56 individuals with thee timepoints. The average number of days between timepoint 1 and 2 was 154.8 ±80.5 (standard deviation, SD) days, and timepoint 2 and 3 was 194.9 ±144.5 (SD) days. To investigate whether there were changes in alpha diversity across time, we performed linear mixed effects models to control for the effect from the individual. Based on the calculations on the longitudinal dataset controlling for the individual, there were no significant increases in the Shannon index, richness, or evenness. Although not significant, there may be an increase in the Shannon index across timepoints 1-3 (Figure 3). Next, we calculated the Bray-Curtis similarity of microbiome composition to investigate changes in the microbiome across time. There was no significant difference from one timepoint to the next (Figure 3), or when comparing the first timepoint with each subsequent timepoint (data not shown). However, there was a shift in the median towards lower Bray-Curtis similarity indices across longitudinal timepoints 1-5 towards a lower similarity index (Figure 3). A permutated multivariate analysis of variance was performed across all timepoints to calculate microbiome variance across longitudinal samples. There was a significant difference between all longitudinal samples from timepoint 1 and timepoint 3 (R^2^ = 0.0088, p = 0.035). Average days separating timepoints 1 and 3 were 335.9 ± 170.5 (SD) days. When computing the variance for subjects with both timepoints, we resolved no significant variation within the microbiome data. Considering the microbiome composition and health and diet survey information, we calculated the microbiome score for each sample and saw a significant increase in the overall microbiome score across timepoints 1-3 (Figure 3).

**Figure 3.**
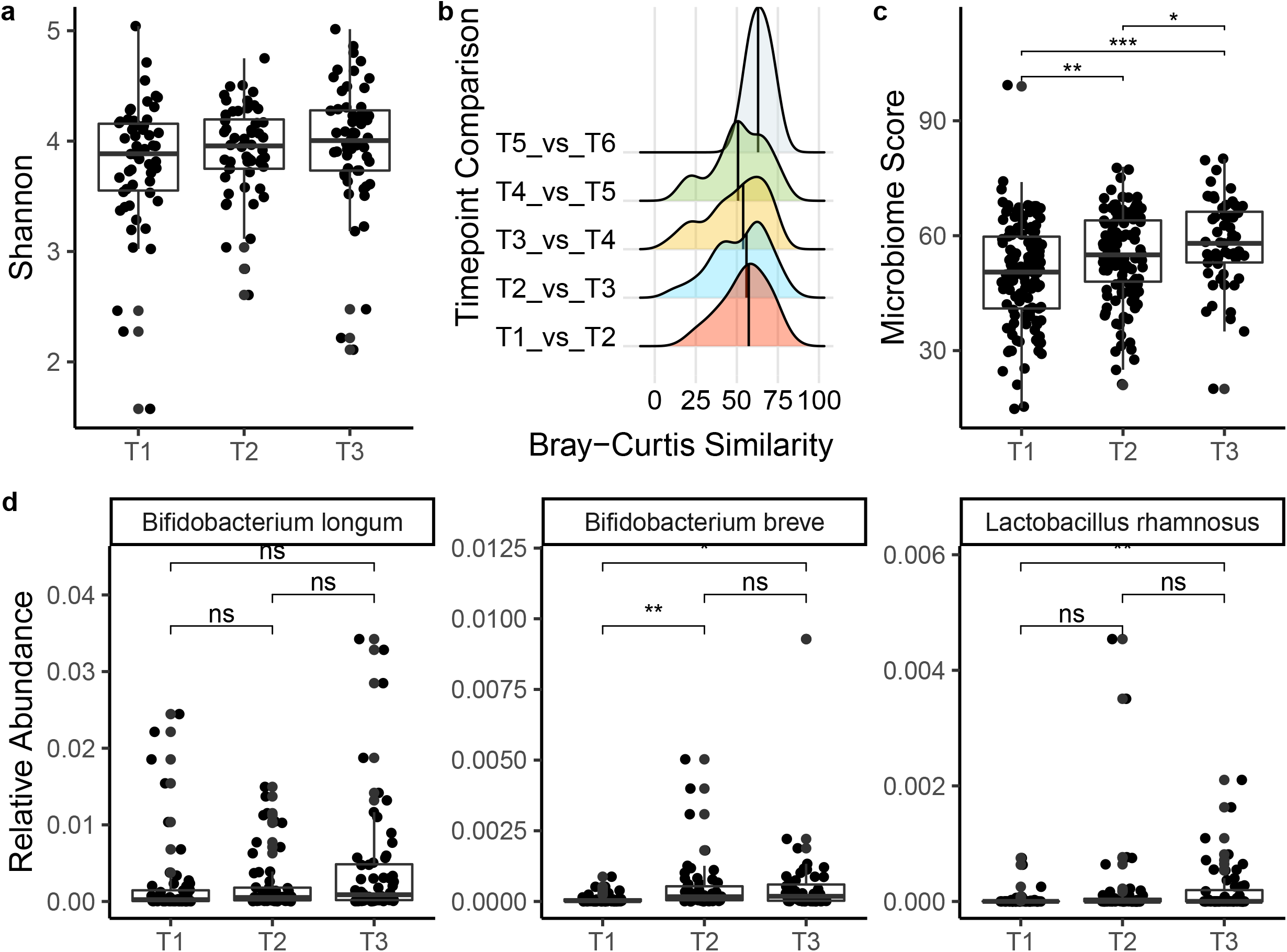
Longitudinal microbiome diversity and relative abundances of probiotics in subjects with IBS. A) Shannon index of the microbiome composition from subjects with timepoints 1 – 3. B) Bray-Curtis similarity of timepoints within each individual. Each timepoint is compared to each subsequent timepoint. C) The overall microbiome score across timepoints 1 – 3. Wilcoxon tests were computed with FDR adjusted p values. D) Relative abundances of probiotic species in subjects across 3 timepoints. T-tests were computed with FDR adjusted p values. * p value < 0.05, ** p value < 0.01, *** p value < 0.001.

Because there were different subtypes associated with IBS included in our population, we investigated whether the individually formulated probiotics targeted towards relieving the symptoms of constipation and diarrhea increased in abundance in the microbiomes of the IBS population. Each formulation contained approximately 4-8 probiotic strains, each at different concentrations. One of the common probiotics formulated for constipation was *Bifidobacterium longum* and the formulations for diarrhea included *Bifidobacterium breve* and *Lactobacillus rhamnosus*. In the longitudinal dataset, *B. breve* and *L. rhamnosus* significantly increased in abundance across time (Figure 3). *B. breve* significantly increased from timepoint 1 to 2 and 3, but there was no significant change between the 2^nd^ and 3^rd^ timepoints (Figure 3). *L. rhamnosus* was significantly increased in abundance at timepoint 3 compared to timepoint 1 (Figure 3). There was not a significant increase in the relative abundance of *B. longum* across timepoints 1-3.

## Discussion

Although IBS is prevalent across the population, the underlying factors contributing to the syndrome makes diagnosis and treatment challenging to define and standardize. Previous amplicon-based studies have identified changes in microbiome composition and diversity in individuals with IBS compared to a healthy control population (29, 30). Concomitant with previous findings, our study corroborates the significant microbial community composition differences and diversity between healthy individuals and people with IBS. Unlike other studies, whole metagenome shotgun sequencing enabled us to identify pathways associated with the dominant subtypes of IBS. In addition, our precision probiotics for individuals with IBS showed a significant microbiome score improvement across time. Clinical studies that administer probiotics to individuals with IBS have shown reduced symptom severity and gut discomfort (25–27). Although we did not find a significant change in alpha or beta diversity in the longitudinal IBS profiles with probiotic supplementation, there was a significant increase in the relative abundances of probiotics detected in the gut microbiomes. Of subjects with three timepoints, 91% had all three of the common probiotic species we included in formulations. These results indicate that probiotic supplementation may be changing microbial community composition and function that may alleviate IBS symptoms. Further research is needed to assess longitudinal changes in microbiome function in response to probiotics in IBS.

In the microbiome composition of individuals with IBS, there was a significant reduction in alpha diversity and anti-inflammatory microbes while there was an increase in inflammatory microbes. A reduction in alpha diversity in the microbiome may indicate a loss of microbial species in response to different environmental factors (i.e., antibiotics) or a presence of microbial players that may be driving the reduction in diversity. For example, there was increased abundance in *Klebsiella* and *Escherichia/Shigella* in small intestinal bacterial overgrowth and a reduced duodenal microbiome diversity (31). Consistent with IBS-A, IBS-C, and Crohn’s disease studies, we found lower relative abundances of the anti-inflammatory microbe *F. prasunitzii* in individuals with IBS than the healthy cohort (29, 32–35). In contrast to previous amplicon-based studies that did not find a reduced abundance of *F. prausnitzii* in IBS-D (35–37), we detected *F. prasunitzii* at lower levels in IBS-D. *F. prausnitzii* enhances gut barrier protection and produces butyrate, a short chain fatty acid essential for gut health (29, 32, 38, 39). *Roseburia intestinalis* has an anti-inflammatory role in the gut and is reduced in individuals with Crohn’s disease (40, 41). *R. intestinalis* was significantly reduced in IBS-C and IBS-D subtype (Figure 2). *Shigella spp*., a major contributor to diarrheal disease (42) and associated with post-infectious IBS (43), was found to be increased in the IBS subtypes (Figure 2). These variations in the microbial signature was taken into account for the internal health index score. The scoring system is highly dependent on the microbial abundance levels in the profile and their association with gastrointestinal conditions like IBS (43, 44) and balance of the gut ecosystem, including the presence *Faecalibacterium* (32–34, 38, 39).

The other differentially abundant microbes have an unclear role in IBS. *Ruminococcus lactaris* is negatively correlated to IL-8 (45) and is more abundant in a non-chronic kidney disease cohort (46), but has also been shown to be associated with a high-fat diet in a murine diabetes model (47). *Eubacterium rectale* is a butyrate producer associated with infant gut microbiome development (48), but is also associated with obesity and dysbiosis (49). In a recent metagenomic assembly study of *E. rectale*, there were different subspecies due to genetic and geographic dispersal in human populations, revealing differences in subspecies physiologies and metabolisms (50). *Prevotella spp*. is common in non-western plant-rich diets (51) and decreased in individuals with constipation (52), but has also been associated with chronic inflammatory conditions (53, 54). These studies indicate that the role of some microbes detected in this study is context and environmentally dependent.

Functional analysis identified pathways associated with each of the phenotypic classifications of IBS. The methanogenesis from an acetate pathway was associated with IBS-C (Figure 2). Methanogenesis contributes to methane production, which is correlated to the severity of constipation (55) and may be useful as a diagnostic indicator of constipation predominant IBS (56, 57). Surprisingly, methanogenesis was also associated with IBS-D. Previous studies have demonstrated the reduction of methanogens in IBS-D (58). However, *Blautia spp*. and *Fusicatenibacter* were microbes detected to have genes that contribute to the methanogenesis pathway (Table S1). *Blautia spp*. and *Fusicatenibacter* produce short-chain fatty acids and gases through carbohydrate fermentation, substrates for methanogenesis. An overabundance of methanogenesis may lead to gut symptoms in IBS. The *Bifidobacterium* shunt was also associated with IBS-C. The *Bifidobacterium* shunt, also called the fructose-6-phosphate shunt, produces short-chain fatty acids (SCFA) and other organic compounds (59, 60). Depending on the chemical and microbial microenvironment, SCFA can regulate the growth and virulence of enteric pathogens (61). In addition, SCFA stimulates water absorption in the colon (62). If too much water is absorbed, the stool becomes more solid, resulting in constipation. Thus, factors affecting host physiology in IBS may depend upon the microenvironments and microbes present in the gut.

The enterobacterial common antigen (ECA) biosynthesis pathway was associated with IBS-A. The ECA is one of the components of the outer membrane of Gram-negative bacteria and its association with IBS-A may indicate the increased presence of *Enterobacterales* in the gut microbiome. Interestingly, the ECA may contribute to virulence and protect enteric pathogens from bile salts and antibiotics (63–65). Bile acids protect the host from infection, contributing to overall gut intestinal health (66). ECA protection against bile acids and antibiotics may make IBS-A challenging to treat with antibiotics and may contribute to dysbiosis. These results suggest that common antibiotic treatments for IBS may not be ideal for alleviating symptoms or treating the possible underlying microbiome triggers associated with IBS-A.

Pathways associated with healthy microbiomes were amino acid and ribonucleotide synthesis, polysaccharide degradation, and fermentation. L-methionine biosynthesis by sulfhydrylation and cysteine biosynthesis implies the presence of hydrogen sulfide in the gut (67). An overabundance of hydrogen sulfide induces inflammation, while low levels protect the gut lining and microbes against reactive oxygen species (68–70). Polysaccharide degradation, specifically beta-mannan degradation, is primarily driven by *Roseburia intestinalis* and the metabolic output has been shown to promote gut health (71). As products of fermentation, the role of SCFA has been implicated in cardiovascular and neurologic pathologies (72–74). The thiamine diphosphate biosynthesis pathway was associated with healthy gut metagenomes while negatively associated with IBS-A (Figure 2). A thiamine deficiency has been shown to increase risk for lifelong neurodevelopmental consequences and is associated with many cardiovascular diseases (75, 76). These results demonstrate the balance of metabolites must be regulated to maintain gut homeostasis and overall health. When the chemical and microbiome balance is disrupted, host physiology may be affected, leading to worsening gut symptoms or onset of disease.

IBS is heterogeneous; a universal cocktail of probiotics may not comprehensively target all symptoms experienced by individuals with IBS. Therefore, one of our goals is to individually formulate prebiotics and probiotics to address the more common symptoms experienced by individuals with IBS. There were three common strains included in formulas to specifically target constipation and diarrhea. *Bifidobacterium longum* was included in formulations for constipation (77, 78). *B. breve* and *Lactobacillus rhamnosus* were included in formulations for diarrhea (79–84). Each of these probiotics were added to formulas for a total of 4-8 different probiotic strains at different concentrations.

In probiotic studies, most strains are detectable for less than 2 weeks following the cessation of probiotic supplementation (85–87). Because individuals in this study took daily probiotic supplements across a longitudinal time period (4 months between each microbiome test), we investigated microbiome changes in response to daily probiotic supplementation. There was no significant change in alpha or beta diversity across time in the population, but there may have been changes in diversity within the individual (Figure 3). However, there was a shift in beta diversity of the microbiome from one timepoint to the next, indicating there may be changes in microbiome composition in response to probiotic supplementation (Figure 3). Thus, diversity metrics that compare population-level information may not show the impact of probiotic usage, but may influence smaller communities within the gut with postbiotic release.

In addition to the diversity metrics calculated, we calculated an overall microbiome score to consider the microbiome composition and health and diet survey. Across timepoints 1-3, there was a significant increase in the microbiome score, indicating an improvement in the overall microbiome and symptoms in response to precision probiotic supplementation. When investigating individual probiotics, *Bifidobacterium longum* did not significantly increase across timepoints in the IBS subjects even when provided in precision formulations. The presence of *B. longum* may promote gut health through cross-feeding mechanisms that lead to the production of short-chain fatty acids (88, 89). *B. breve* and *L. rhamnosus* increased in relative abundance across time in individuals with IBS, indicating colonization of the gut microbiome that may contribute to positive changes in microbiome physiology. Further investigation is needed to identify potential functional changes in microbiome metabolism with daily prebiotic and probiotic supplementation in IBS and whether symptoms associated with IBS can be improved.

There are several limitations to this current study. First, the self-reporting nature of IBS is a limitation to this study. For official diagnosis of IBS, the Rome IV criteria assesses symptoms related to stool consistency and appearance, recurrent abdominal pain, and changes in bowel habits (90, 91). Although the health and diet questionnaire included questions regarding gut symptoms and chronic conditions, a formal diagnosis was not verified. For potential life-style modifications in addition to probiotic supplementation, diet changes may also be an important factor in alleviating symptoms or changing the microbiome (92–94). Low FODMAP diet (LFD) and low lactose diet (LLD) reduced the IBS-SSS score.

IBS subjects on LFD had significantly less abdominal pain, bloating, and gas production (93). These diet interventions were not accessed in this study. Second, this study was not designed to investigate longitudinal assessments of comprehensive gut issues experienced by the individuals with IBS. This hindered us to identify whether gut symptoms were alleviated by daily probiotic supplementation or whether there were associations with certain probiotic formulations in improving certain symptoms in IBS. However, because the relative abundance of the common probiotics formulated for constipation and diarrhea were increased in relative abundance across time, these results may inform future studies. Additional research is also needed to determine the roles of specific pathways in the etiology of IBS.

In summary, we reported differentially abundant microbes and functional pathways associated with IBS. We also identified an increased relative abundance of probiotics in the gut microbiomes of people with IBS across time. These data may help inform future studies and therapeutic strategies by identifying important microbes and pathways associated with each IBS subtype. As IBS is a multi-factorial syndrome, there is no one-size-fits-all approach to target all symptoms experienced by individuals with IBS. A combination of diet and probiotics may be needed to alleviate symptoms of IBS. Longitudinal monitoring of the gut microbiome is also important to understand changes associated with symptom progression. Precision probiotics may help target individual needs, although further research is needed to identify the pathway benefits of prebiotic and probiotic supplementation on health.

## Materials and Methods

### Participants and sample collection

Users of our (Sun Genomics, San Diego) commercial gut microbiome testing component (Floré Gut Health Test Kit) submitted a stool sample for metagenomic sequencing. The stool sample was collected by the user with provided gut testing kit instructions. Samples were collected in accordance with IRB # SG-04142018-001 with informed consent form 001-B. A sterile swab was used for the first collection device to collect and store the stool sample in a collection tube with stabilization buffer. The second sample was collected via the Easi-Collect device (GE). Samples were mailed via FedEx to the Sun Genomics lab for analysis.

A total of 611 participants were included in this study (Table 1). All participants completed a health and diet survey that asked questions about health status and dietary preferences. The control population included in this study was self-reported as healthy with no listed comorbidities with a BMI range from 18.5 – 25 (Table 1) (95). The IBS population was also self-reported and included the symptoms associated with the syndrome, including constipation, diarrhea, a mix of both constipation and diarrhea, or unspecified.

### Metagenomic sequencing and analysis

For DNA extractions, samples were first processed with a tissue homogenizer and then lysed with a lysis buffer and proteinase K. DNA was extracted and purified with a proprietary method (Patent #10428370 and #10837046 - Universal Method For Extracting Nucleic Acids Molecules From A Diverse Population Of One Or More Types Of Microbes In A Sample). Library preparation was performed with DNA sonication, end-repair, and adaptor ligation with NEBNext reagents. Size selection was performed with MagJet Magnetic Beads according to manufacturer instructions. Library quantitation was performed with qPCR and sequenced on an Illumina NextSeq 550 (Illumina, San Diego). After sequencing, reads were quality filtered and processed. Metagenomic reads were decontaminated from human reads using Bowtie2. After decontamination, there was an average of 6,581,844 reads per sample (SD = 4,426,117) with a minimum of one million reads to be included in downstream analyses. Next, reads were aligned to a hand curated database of over 23,000 species. Humann3 was used for pathway analysis (96). Pathway abundance was normalized to copies per million (cpm).

### Statistical analyses

All statistical analyses were performed in R. Principal coordinates analysis was performed with a Bray-Curtis dissimilarity matrix to compare between sample diversities. Within sample diversity was calculated with the Shannon diversity index. To calculate variance between samples based on metadata classifications, permutational multivariate analysis of variance (PERMANOVA) was performed with the “adonis” function from the “vegan” package (97). Specifically, the influence of health status was computed across the microbiome composition and pathway abundance profiles. MaasLin2 was used for distinguishing pathway features between healthy and IBS subtypes (98).

